# The two-component system CroRS regulates isoprenoid flux to mediate antimicrobial tolerance in the bacterial pathogen *Enterococcus faecalis*

**DOI:** 10.1101/2022.12.05.519242

**Authors:** Francesca O Todd Rose, Rachel L Darnell, Sali Morris, Olivia Paxie, Georgia Campbell, Gregory M Cook, Susanne Gebhard

## Abstract

Antimicrobial tolerance is the ability of a microbial population to survive, but not proliferate, during antimicrobial exposure. Significantly, it has been shown to precede the development of bona fide antimicrobial resistance. We have previously identified the two-component system CroRS as a critical regulator of tolerance to antimicrobials like teixobactin in the bacterial pathogen *Enterococcus faecalis*. To understand the molecular mechanism of this tolerance, we carried out RNA-seq analyses in the *E. faecalis* wild-type and isogenic Δ*croRS* mutant to determine the teixobactin-induced CroRS regulon. We identified a 132 gene CroRS regulon and show CroRS upregulates expression of all major components of the enterococcal cell envelope in response to teixobactin challenge. To gain further insight into the function of this regulon we isolated and characterized Δ*croRS* mutants recovered for wild-type growth and tolerance. We show introduction of a single stop codon in a heptaprenyl diphosphate synthase (*hppS*), a key enzyme in the synthesis of the quinone electron carrier demethylmenaquinone (DMK), is sufficient to rescue loss of cell envelope integrity in the *croRS* deletion strain. Based on these findings, we propose a model where CroRS acts as a gate-keeper of isoprenoid biosynthesis, mediating flux of isoprenoids needed for cell wall synthesis (undecaprenyl pyrophosphate; UPP) and respiration (DMK) to maintain cell wall homeostasis upon antimicrobial challenge. Dysregulation of this flux in the absence of *croRS* leads to a loss of tolerance, which is rescued by loss of function mutations in HppS, allowing an increase in isoprenoid flow to UPP and subsequently cell wall synthesis.

**Importance:** Antimicrobial tolerance is the ability of a microorganism to survive, but not grow upon antimicrobial challenge, and is an important precursor to the development of antimicrobial resistance (the ability to profilerate). Understanding the molecular mechanisms that underpin tolerance will therefore aid in hampering the development of resistance to novel antimicrobials such as teixobactin. CroRS is an essential two-component regulator of antimicrobial tolerance in the bacterial pathogen *Enterococcus faecalis*. We have determined the antimicrobial-induced CroRS regulon and identified key mutations in a heptaprenyl diphosphate synthase to uncover a novel mechanism of antimicrobial tolerance.

## Introduction

The emergence of multidrug-resistant bacterial pathogens has rendered standard treatments ineffective, allowing infections to persist and spread. Significantly, antimicrobial tolerance (AMT), i.e., the ability of a bacterium to survive but not proliferate during antimicrobial exposure, has been shown to precede the development of bona fide antimicrobial resistance (1–4). Enterococci are a group of Gram-positive bacteria that inhabit a wide variety of ecological niches such as food, fresh water and the gastrointestinal tract of humans, animals and insects (5). Although primarily commensals, enterococci are also clinically significant opportunistic pathogens that can exploit a compromised host to cause diseases such as urinary tract infections, bacteremia, and endocarditis (6). *Enterococcus faecalis* and *Enterococcus faecium* are the most abundant enterococcal species in humans and a leading cause of hospital-acquired infection (7).

Two-component systems compose an important regulatory network of the cell envelope stress response in *Enterococcus faecalis*, with a number implicated in antimicrobial resistance i.e., vancomycin resistance, VanRS, and cephalosporin resistance, CroRS (8–11). In addition, we have previously identified CroRS as a critical regulator of antimicrobial tolerance (12). CroRS is encoded on a bicstronic operon with *croS* encoding the sensor kinase and *croR* its cognate response regulator (8). Taken together, these studies demonstrate the crucial role of CroRS in mediating the response to antimicrobial attack in *E. faecalis*. However, the precise architecture ‘ of the response network and the molecular mechanism(s) conferring this tolerance require further elucidation.

Previous transcriptional analyses of the CroRS regulon have used RNA-Seq to identify genes differentially expressed in the presence and absence of antimicrobial stress (11, 13). Muller *et al* (2018) identified 50 potential CroR-regulated genes in the *E. faecalis* JH2-2 strain in the absence of antimicrobial stress (11), while Timmler *et al* identified 87 CroR-regulated genes differentially expressed in the *E. faecalis* OG1RF strain in the presence of bacitracin-induced antimicrobial stress (Timmler *et al.*, 2022). In the OG1RF strain CroS has two cognate response regulators CroR and CisR, of which CisR is notably absent in the JH2-2 strain (14). In addition, while CroRS is known to respond to the presence of bacitracin, *E. faecalis* displays low tolerance to bacitracin (12, 13). Therefore, the tolerance-inducing CroRS regulon remains undiscovered.

Teixobactin (TXB) represents a new class of antimicrobial with a unique chemical scaffold and lack of detectable resistance (15). It has proven efficacy against multidrug-resistant pathogens such as enterococci, staphylococci and *Mycobacterium tuberculosis* and has been shown to lo induce cell lysis through binding of cell wall precursors lipid II and lipid III in *Staphylococcus aureus* (15, 16). Further to this mechanism, recent studies suggest that TXB potency is amplified by the formation of TXB-lipid II clusters (17, 18). The unique enduracididine C-terminal headgroup of TXB specifically binds to the conserved pyrophosphate-saccharide moiety of cell wall precursors such as lipid II and lipid III while the N-terminus coordinates with a second pyrophosphate from another lipid II molecule (18). Clustering of lipid II within this structure is thought to displace phospholipids and disrupt the membrane, generating a simultaneous action against cell wall synthesis and the cytoplasmic membrane to produce a highly effective antimicrobial (18).

Antimicrobials are thought to kill bacteria through interaction with specific intracellular targets (19). Antimicrobial drug-target interactions, and their respective direct effects, are generally well characterized. By contrast the bacterial responses to antimicrobial treatments that contribute to cell death are not as well understood and have proven to be complex as they involve many genetic and biochemical pathways (19). It is currently unknown how CroRS protects against killing by cell wall-targeting antimicrobials such as TXB and the glycopeptide vancomycin. However, by determining the antimicrobial-induced CroRS regulon, we can gain a better understanding of the physiological processes this regulatory system controls, and begin to uncover its role in AMT. In this study we determine the TXB-induced CroRS regulon and identify key genes and pathways involved in mediating CroRS-regulated AMT. We show CroRS regulates expression of all major pathways of cell envelope biosynthesis and functions as a gatekeeper of isoprenoid flux between cell wall biosynthesis and respiratory energy metabolism to confer AMT in *E. faecalis*.

## Results and Discussion

### Whole-genome transcription profiling of the *E. faecalis* WT and Δ*croRS* mutant in the presence and absence of teixobactin

To understand how CroRS contributes to TXB tolerance, we first aimed to identify which genes it controls in response to TXB exposure. Previous optimisation with the *E. faecalis* JH2-2 wild-type (WT) showed challenge with 0.5 *μ*g ml^-1^ of TXB for 1 h on mid-exponential phase cells was optimal for inducing a CroRS response without significantly impacting growth (12). These conditions were also deemed appropriate for the Δ*croRS* strain, with no difference in growth inhibition observed between Δ*croRS* and the WT under these conditions (**Figure S1**) (12). To identify the TXB-induced CroRS regulon, four different gene expression profiles (> 1.0 fold-log_2_) were generated following RNA sequencing: (1) WT untreated vs Δ*croRS* untreated (**Table S1**), (2) WT treated vs Δ*croRS* treated (**Table S2**), (3) WT treated vs untreated (**Table S3**), and (4) Δ*croRS* treated vs untreated (**Table S4**). These four gene expression profiles were then fed into a filtering system to isolate the TXB-induced CroRS regulon (**Figure S2**). This filtering system was deliberately chosen for its stringence to allow us to specifically identify genes controlled by CroRS in response to TXB and their role in TXB tolerance.

#### The teixobactin-induced CroRS regulon

A total of 538 genes were differentially expressed (>1.0 fold-log_2_) in the WT vs Δ*croRS* in the presence of TXB (**Table S1**). However, only 132 of these genes were found to belong to the TXB-induced CroRS regulon (**Table S5**). Of these 132 genes, 117 were upregulated and 15 were downregulated. To identify metabolic pathways up- and downregulated by CroRS in response to TXB challenge, genes were categorized into KEGG gene ontologies using the well-defined *E. faecalis* V583 strain as a reference.

Over 25% of genes upregulated by CroRS were involved in cell envelope biogenesis (**Figure 1A**). Strikingly, this involved all layers of the enterococcal cell envelope, i.e., the cytoplasmic membrane (lipid metabolism), the peptidoglycan cell wall (metabolism of terpenoids and polyketides and drug resistance), teichoic acids and cell wall polysaccharides (glycan biosynthesis and metabolism) (**Figure 1A; Figure 2**). Other pathways highly upregulated included amino acid biosynthesis and membrane transport (**Figure 1A**). In comparison, carbon metabolism, membrane transport and signal transduction were downregulated by CroRS in response to TXB (**Figure 1C**). In addition hypergeometric testing confirmed significant upregulation of cell envelope biogenesis, and the biosynthesis and metabolism of amino acids (**Figure 1B**), as well as downregulation of heme transport, oxidative phosphorylation, and signal transduction (**Figure 1D**).

**Figure 1.**
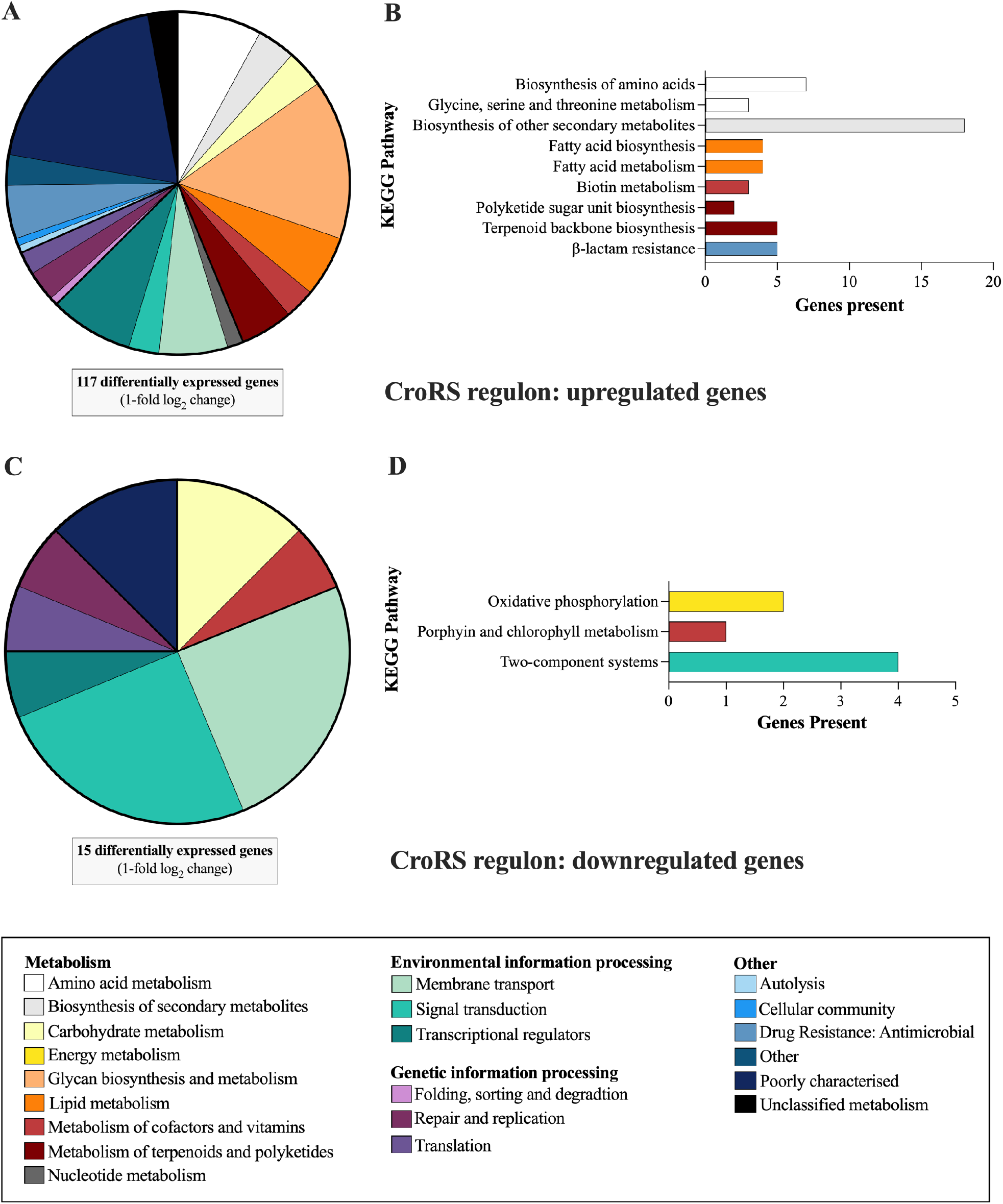
Functional classification and distribution of TXB-induced CroRS-regulated genes. Genes differentially expressed (>1-fold log_2_) in the TXB-induced CroRS regulon were assigned to the well-defined *E. faecalis* V583 KEGG ontologies. These pie charts represent the distribution of these ontologies up and downregulated by the TXB-induced CroRS regulon (A and C). In addition, hypergeometric testing was performed on the TXB-induced CroRS regulon to identify ontologies significantly (P= <0.05) up and downregulated (B and D).

**Figure 2.**
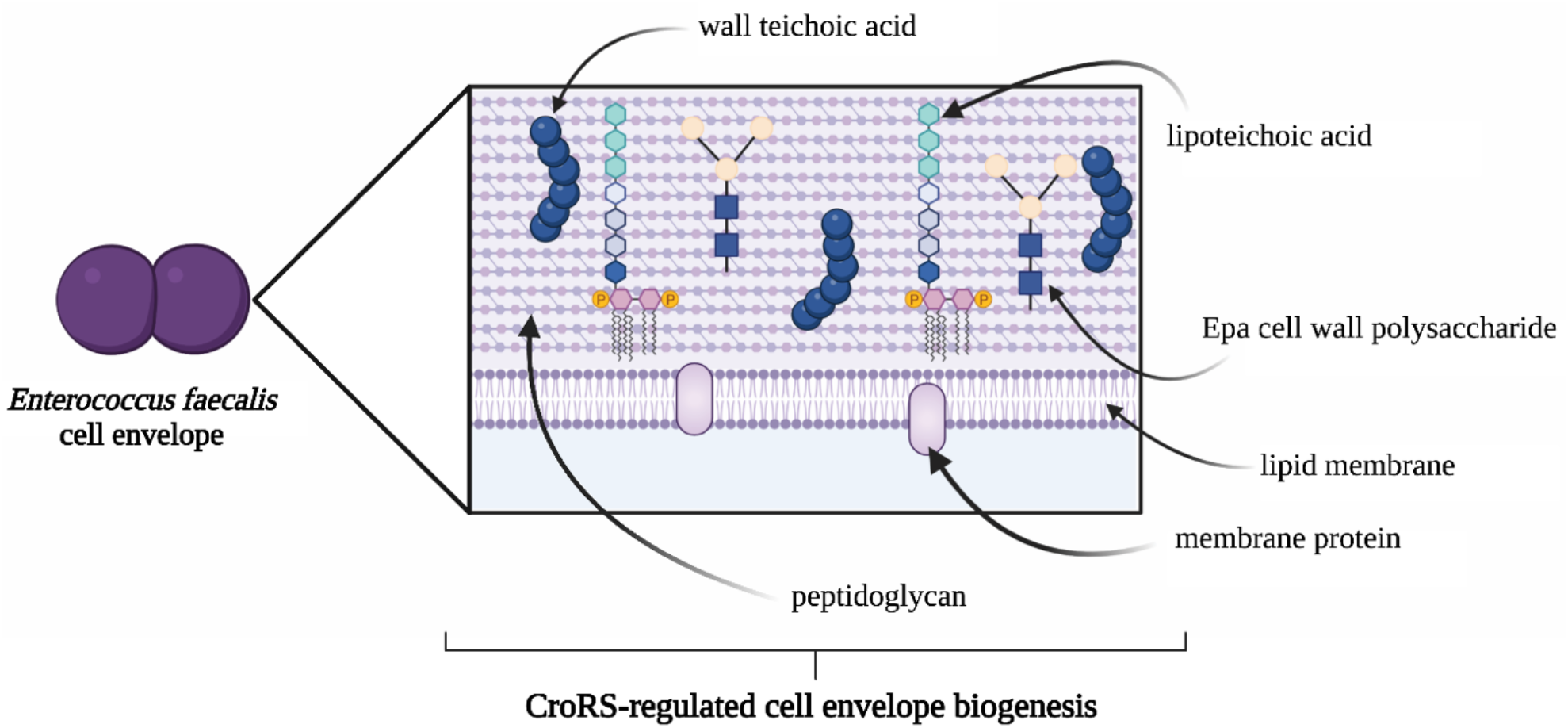
CroRS-regulated pathways of the enterococcal cell envelope in response to TXB stress. The enterococcal cell envelope is composed of two main layers: a cytoplasmic lipid membrane, surrounded by a thick cell wall. Peptidoglycan is the major component of the cell wall, with teichoic acids (wall and lipoteichoic) and the Epa (enterococcal polysaccharide antigen) cell wall polysaccharides as the two other major constituents. CroRS regulates expression of genes required for the biosynthesis of each of these components in response to teixobactin challenge.

### CroRS regulates all aspects of cell envelope biogenesis

CroRS was found to upregulate genes involved in the biogenesis of all major components in the enterococcal cell envelope in response to TXB challenge (**Table 1; Figure 2**). To begin, enterococci utilise the mevalonate (MVA) pathway to synthesize the isopenoid precursors farnesyl pyrophosphate (FPP) and isoprenyl pyrophosphate (IPP) for generation of the essential cell wall lipid carrier undecaprenyl pyrophosphate (UPP), and quinones such as demethylmenaquinone (DMK) for electron transport. In enterococci, six genes make up the MVA pathway, five of which were upregulated (2.9 – 4.6 fold-log_2_) by CroRS in response to TXB (**Table 1**). The sixth, *mvk* (EF0904) looks to be in an operon with *mvaD* and *mvaK*, and fulfills all criteria for the CroRS regulon except for its observed downregulation (−2.1 fold-log_2_) in the WT versus Δ*croRS* strain in the absence of TXB (**Table S1**). It was also shown to be differentially expressed in the presence versus absence of CroR by Muller *et al* (11). Therefore, we hypothesize that *mvk* is regulated by CroRS, but its expression may also be controlled by a secondary regulator.

**Table 1.**
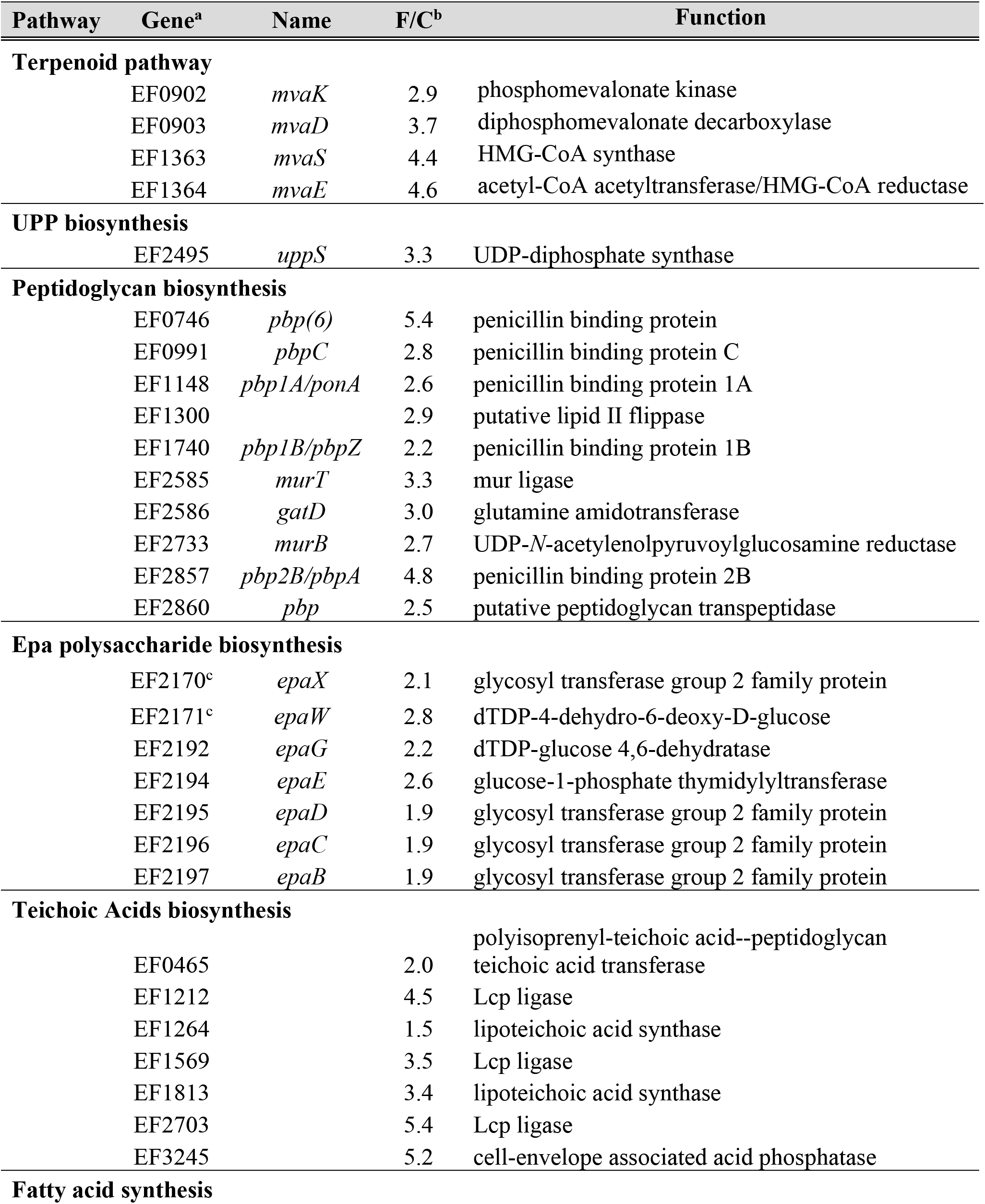
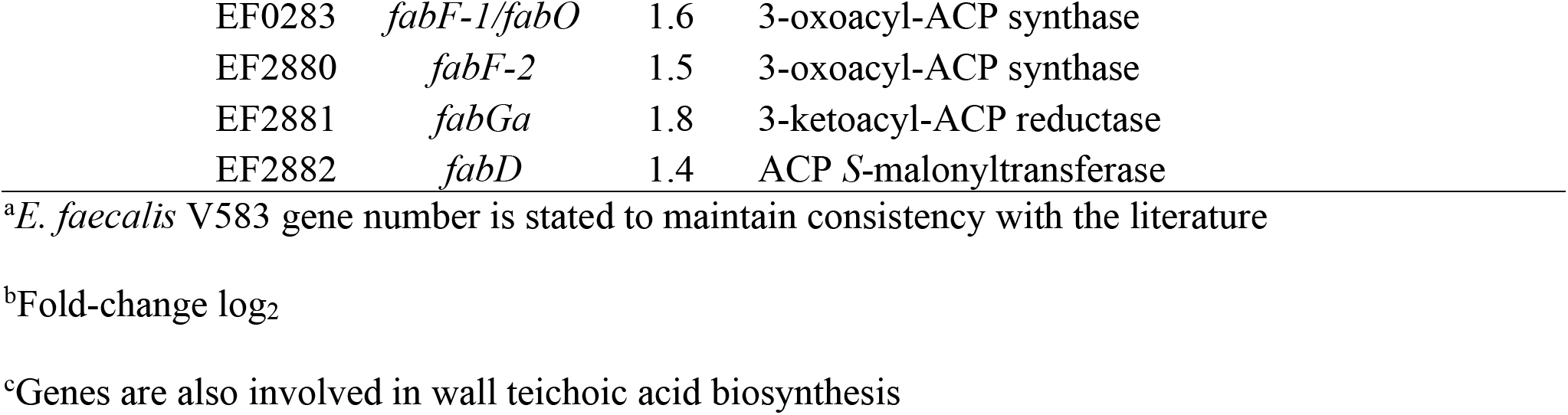
Differential gene expression in the *E. faecalis* WT vs Δ*croRS* strain in the presence of TXB.

| Pathway | Gene<sup>a</sup> | Name | F/C<sup>b</sup> | Function |
| --- | --- | --- | --- | --- |
| **Terpenoid pathway** |  |  |  |  |
|  | EF0902 | *mvaK* | 2.9 | phosphomevalonate kinase |
|  | EF0903 | *mvaD* | 3.7 | diphosphomevalonate decarboxylase |
|  | EF1363 | *mvaS* | 4.4 | HMG-CoA synthase |
|  | EF1364 | *mvaE* | 4.6 | acetyl-CoA acetyltransferase/HMG-CoA reductase |
| **UPP biosynthesis** |  |  |  |  |
|  | EF2495 | *uppS* | 3.3 | UDP-diphosphate synthase |
| **Peptidoglycan biosynthesis** |  |  |  |  |
|  | EF0746 | *pbp(6)* | 5.4 | penicillin binding protein |
|  | EF0991 | *pbpC* | 2.8 | penicillin binding protein C |
|  | EF1148 | *pbp1A/ponA* | 2.6 | penicillin binding protein 1A |
|  | EF1300 |  | 2.9 | putative lipid II flippase |
|  | EF1740 | *pbp1B/pbpZ* | 2.2 | penicillin binding protein 1B |
|  | EF2585 | *murT* | 3.3 | mur ligase |
|  | EF2586 | *gatD* | 3.0 | glutamine amidotransferase |
|  | EF2733 | *murB* | 2.7 | UDP-*N*-acetylenolpyruvoylglucosamine reductase |
|  | EF2857 | *pbp2B/pbpA* | 4.8 | penicillin binding protein 2B |
|  | EF2860 | *pbp* | 2.5 | putative peptidoglycan transpeptidase |
| **Epa polysaccharide biosynthesis** |  |  |  |  |
|  | EF2170<sup>c</sup> | *epaX* | 2.1 | glycosyl transferase group 2 family protein |
|  | EF2171<sup>c</sup> | *epaW* | 2.8 | dTDP-4-dehydro-6-deoxy-D-glucose |
|  | EF2192 | *epaG* | 2.2 | dTDP-glucose 4,6-dehydratase |
|  | EF2194 | *epaE* | 2.6 | glucose-1-phosphate thymidyltransferase |
|  | EF2195 | *epaD* | 1.9 | glycosyl transferase group 2 family protein |
|  | EF2196 | *epaC* | 1.9 | glycosyl transferase group 2 family protein |
|  | EF2197 | *epaB* | 1.9 | glycosyl transferase group 2 family protein |
| **Teichoic Acids biosynthesis** |  |  |  |  |
|  | EF0465 |  | 2.0 | polyisoprenyl-teichoic acid--peptidoglycan teichoic acid transferase |
|  | EF1212 |  | 4.5 | Lcp ligase |
|  | EF1264 |  | 1.5 | lipoteichoic acid synthase |
|  | EF1569 |  | 3.5 | Lcp ligase |
|  | EF1813 |  | 3.4 | lipoteichoic acid synthase |
|  | EF2703 |  | 5.4 | Lcp ligase |
|  | EF3245 |  | 5.2 | cell-envelope associated acid phosphatase |
| **Fatty acid synthesis** |  |  |  |  |

|  |  |  |  |
| --- | --- | --- | --- |
| EF0283 | *fabF-1/fabO* | 1.6 | 3-oxoacyl-ACP synthase |
| EF2880 | *fabF-2* | 1.5 | 3-oxoacyl-ACP synthase |
| EF2881 | *fabGa* | 1.8 | 3-ketoacyl-ACP reductase |
| EF2882 | *fabD* | 1.4 | ACP *S*-malonyltransferase |

<sup>a</sup>*E. faecalis* V583 gene number is stated to maintain consistency with the literature
<sup>b</sup>Fold-change log<sub>2</sub>
<sup>c</sup>Genes are also involved in wall teichoic acid biosynthesis

Following synthesis and translocation across the membrane, the peptidoglycan precursor lipid II is integrated into the peptidoglycan matrix by penicillin-binding proteins.We observed an upregulated expression of six (of a total eight) penicillin binding proteins (pbp) by CroRS in response to TXB (**Table 1**). This included the transglycosylases *pbp1A* and *pbp1B*, the transpeptidase *pbp2B*, the unclassified *pbpC* and *pbp(6)*, and the putative peptidoglycan transpeptidase EF2860 (**Table 1**). CroRS and penicillin binding proteins PBP2B and PBP1A have previously been associated with cephalosporin resistance in *E. faecalis* and *E. faecium* (20–23). This suggests CroRS-mediated cephalosporin resistance may occur through regulation of these penicillin binding protein genes. In addition, PBPC and PBP2B have also been identified as critical and important, respectively, for growth of *E. faecalis*, further highlighting the crucial role of CroRS target genes in cell function (21).

The enterococcal polysaccharide antigen (Epa) is a major component of the enterococcal cell wall. Epa biosynthesis is encoded by a locus comprised of two gene clusters, one conserved and one variable (24–27). The conserved gene cluster (EF2198-EF2177; *epaA - epaR*) is responsible for the biosynthesis of the rhamnopolysaccharide backbone and mutations in *epaB* and *epaE* have been shown to increase susceptibility to salt and cell stress (28, 29); while the variable gene cluster (EF2177-EF2164) is responsible for the biosynthesis and assembly of the teichoic acids covalently linked to the backbone (26, 28). This variable region represents the major differences in Epa between *E. faecalis* isolates and genes in this cluster, such as the *epaX-like* gene (EF2170), have been shown to reduce peptidoglycan crosslinking and resistance to the autolytic compound lysozyme. In addition, mutations loss of Epa has been associated with altered susceptibility to antimicrobials (29–31). Here, we showed CroRS regulates expression of genes in the conserved gene cluster (*epaB, C, D, E*, and *G*), as well as two variable genes *epaW* and *epaX* (**Table 1**). Therefore, we hypothesize CroRS-regulation of these Epa genes may influence tolerance to TXB-induced cell lysis.

### CroRS regulates expression of the cytochrome *bd* oxidase and NADH oxidase in response to antimicrobial stress

Cytochrome *bd* (*cydA* and *cydB*) is a terminal respiratory oxidase in the electron transport chain that accepts electrons from DMK. It requires heme as cofactor to convert O_2_ to H_2_O and cycle hydrogen to generate a proton motive force and allow the generation of ATP. Cytochrome *bd* can enhance bacterial resistance to oxidative stress, likely through the reduction of intracellular O_2_, and thus potential for ROS formation (32); and CroRS itself is induced by H_2_O_2_ (33). However, CroRS was seen to downregulate expression of cytochrome *bd* (*cydA* −1.8 and *cydB*-1.6-fold log_2_) in response to TXB (**Table S5**) (34). The rationale for this is intriguing as oxidative stress has been proposed as a central, albeit controversial, mechanism of bactericidal antimicrobials (35, 36). Given this, we hypothesize CroRS may function to inhibit aerobic respiration and prevent the generation of endogenous hydroxyl radicals to confer AMT.

### Experimental evolution of *E. faecalis* Δ*croRS* for growth recovery

In concert with the RNAseq analyses, we aimed to identify focal pathways controlled by CroRS through isolation of suppressor mutations via serial passaging experiments. Initial attempts to rescue the Δ*croRS* strain for tolerance were carried out in the presence of TXB, however this failed to produce suppressor mutants able to grow above the Δ*croRS* MIC for TXB. To reduce the selection pressure applied during the experiment we instead exploited the known growth defect of Δ*croRS*, which shows a reduced growth rate compared to the wild-type (10, 37). We hypothesized that evolving the deletion strain for restoration of fast growth may result in suppressor mutants that can correct the physiological defect caused by the loss of CroRS, and that this may in turn also restore AMT. In brief, *E. faecalis* Δ*croRS* was serially passaged into fresh liquid medium every two days to select for faster growing mutants in the population. High turbidity of an overnight growth culture was indicative of restored WT-like growth behaviour. After 10 – 14 days, five clones, one from each independently evolved line, were isolated for further characterisation and named 1BS – 5BS. Growth curves of these mutants demonstrated a decrease in both the lag and exponential doubling time compared to the Δ*croRS* parent strain (**Figure S3)**.

CroRS is activated in the presence of a number of cell-wall acting antimicrobials (12, 13, 38). To determine whether the growth-passaged (BS) mutants were also recovered for AMT, antimicrobial susceptibility assays were carried out for the CroRS-activating cell wall antimicrobials, vancomycin, TXB and bacitracin. Despite evolution in the absence of drugs, all of the growth-passage (BS) mutants were rescued for tolerance, as determined by MBC (minimum bactericidal concentration) values, to TXB and vancomycin but experienced a decrease in resistance, as determined by MIC values (**Table 2**). This indicates that resistance and tolerance are mechanistically very different, with tolerance intricately linked to cell physiology.

**Table 2.** Antimicrobial susceptibility and whole-genome sequence analysis of the evolved Δ*croRS* mutants.

| Strain | Vancomycin |  | Teixobactin |  | Bacitracin |  | Gene | SNP | Codon |
| --- | --- | --- | --- | --- | --- | --- | --- | --- | --- |
|  | MIC | MBC | MIC | MBC | MIC | MBC |  |  |  |
| WT | 1 | >128 | 2 | 16 - 32 | 16 | 64 |  |  |  |
| *ΔcroRS* | 0.25 | <4 | 2 | <4 | 8 - 16 | 8 - 16 |  |  |  |
| 1BS | 0.125 | 128 | 1 | 8 - 16 | 16 | 32 | heptaprenyl diphosphate synthase; *hppS* (EF2057) | 216_225del | K72 |
|  |  |  |  |  |  |  | *N*-acetylmuramoyl-L-amidase; (EF2367) | 52C>T | Q18* |
| 2BS | 0.125 | 128 | 1 | 16 | 16 | 32 | heptaprenyl diphosphate synthase; *hppS* (EF2057) | 567T>A | Y189* |
| 3BS | 0.125 | 128 | 1-2 | 16 | 16 | 32 | DNA-directed RNA polymerase subunit delta; *rpoE* (EF1146) | 584_585delTTinsAA | V195E |
|  |  |  |  |  |  |  | hypothetical protein; (EF0062) | 3848A>T | K1283I |
| 4BS | 0.125 | 128 | 1 | 32 | 16 | 32 | heptaprenyl diphosphate synthase; *hppS* (EF2057) | 91G>T | E31* |
| 5BS | 0.125 | 128 | 1 | 32 | 32 | 32 | phosphate transporter ATP-binding protein; *pstBB* (EF1756) | 661A>G | T221A |
|  |  |  |  |  |  |  | *N*-acetylmuramoyl-L-amidase; (EF2367) | 482T>G | V161G |
|  |  |  |  |  |  |  | hypothetical protein; (EF0062) | 3848A>T | K1283I |

2 MIC and MBCs are given as the median range in $\mu\text{g ml}^{-1}$ and are representative of at least biological triplicate.

Whole genome sequencing of these mutants revealed key mutations in a heptaprenyl diphosphate synthase, *hppS* (EF2057;1BS, 2BS, 4BS), as well as the delta subunit of the DNA-directed RNA polymerase, *rpoE* (EF1146; 3BS), an *N*-acetylmuramoyl-L-amidase (EF2367; 1BS, 5BS), a hypothetical protein (EF0062; 3BS, 5BS), and *pstBB*, a phosphate ABC transporter ATP-binding protein (EF1756; 5BS) (**Table 2**). HppS is a key enzyme in the biosynthesis of DMK, the quinone redox carrier for electron transport, and catalyzes the formation of heptaprenyl pyrophosphate (HepPP) from FPP and IPP (39) (**Figure 5**). HepPP is then used as a substrate by the enzyme MenA (1,4-dihydroxy-2-napthoate octaprenyl transferase) to catalyze the formation of DMK. In two of the three strains carrying *hppS* mutations, 2BS and 4BS, it was the only mutation identified (**Figure S4**), providing strong evidence for HppS as the causative agent for the rescued phenotype. This apparent central role of HppS in the physiology of Δ*croRS* correlates with our observations from the RNA-Seq experiments which showed CroRS-mediated induction of the mevalonate pathway in response to TXB and may imply that CroRS has effects on cell wall biosynthesis also in the absence of antimicrobial challenge.

**Figure 3.**
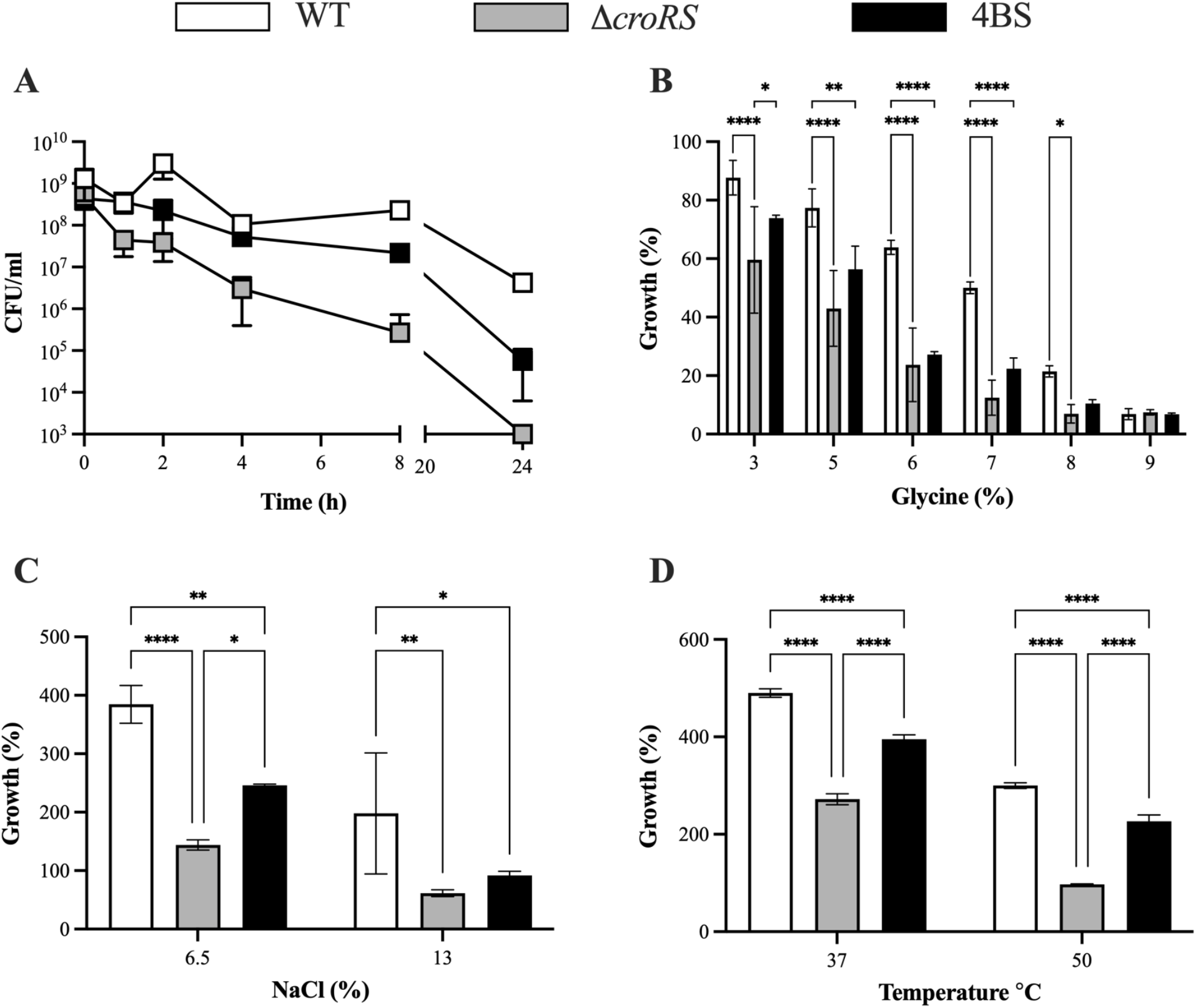
Phenotypic characterization of the WT, Δ*croRS* and 4BS strains. Cell stress assays were used to characterize the *E. faecalis* JH2-2 WT (white), Δ*croRS* (grey), and 4BS mutant (black). In the TXB time-kill assay Strains were grown to mid-exponential phase and challenged with teixobactin (25 × MIC) for 24 h. Cell survival was determined at time = 1, 2, 4, 8, and 24 h post-challenge (**A**). (**B**) Overnight cultures were diluted 1/400 and challenged with a range of glycine concentrations (0, 3, 5, 6, 7, 8, 9%). Growth (%) was determined after 24 h and normalised to an untreated control. (**C** and **D**) Overnight cultures were were diluted to an OD_600_ of 0.05 and challenged with osmotic (NaCl; 6.5% and 13%) (**C**) and temperature stress (37°C and 50°C) (**D**). Growth (%) was determined after 24 h and normalised to an untreated control. All experiments were carried out in at least biological triplicate and are presented as the mean ± SD. A two-way ANOVA was used to determine statistical significance P= <0.05 (* <0.05, *** <0.001, **** <0.0001).

**Figure 4.**
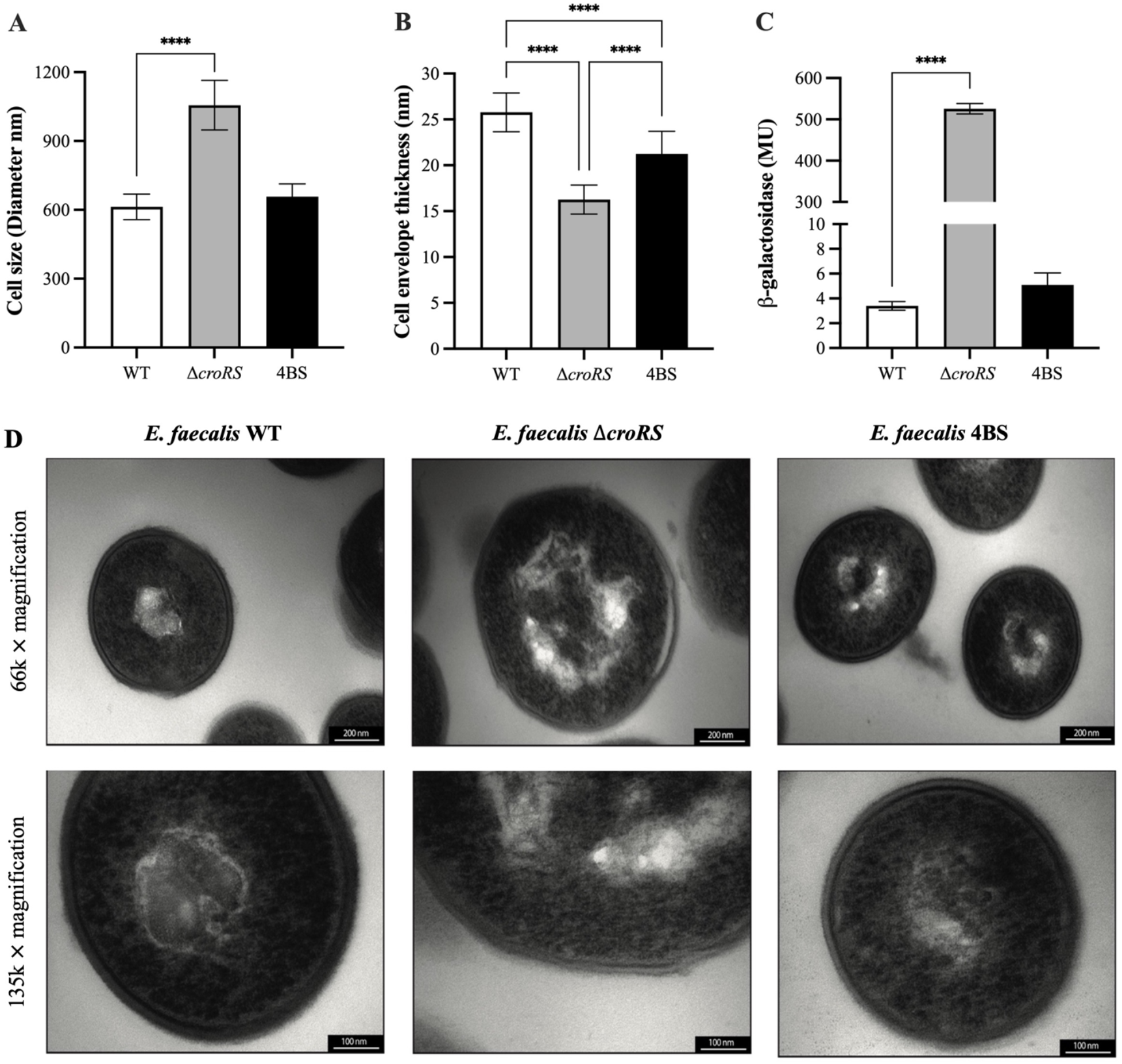
Analysis of the cell envelope stress response in the *E. faecalis* WT, Δ*croRS* and 4BS strain using transmission electron microscopy and the PliaX reporter construct. Strains were grown to mid-exponential phase under normal growth conditions and processed for transmission electron microscopy (TEM) (**D**). TEM images were captured at 66,000 × or 135,000 × magnification. Following TEM imaging a selection of at least twenty cells from individual images were analysed for cell size (**A**) and cell envelope thickness (**B**) using ImageJ. The promoter region of LiaX (regulated by LiaFSR) was fused to *lacZ* and introduced into all three strains. Each strain was then assayed for *β*-galactosidase activity, expressed in Miller units (MU) (**C**). A one-way ANOVA was used to determine statistical significance; P= **** <0.001.

**Figure 5.**
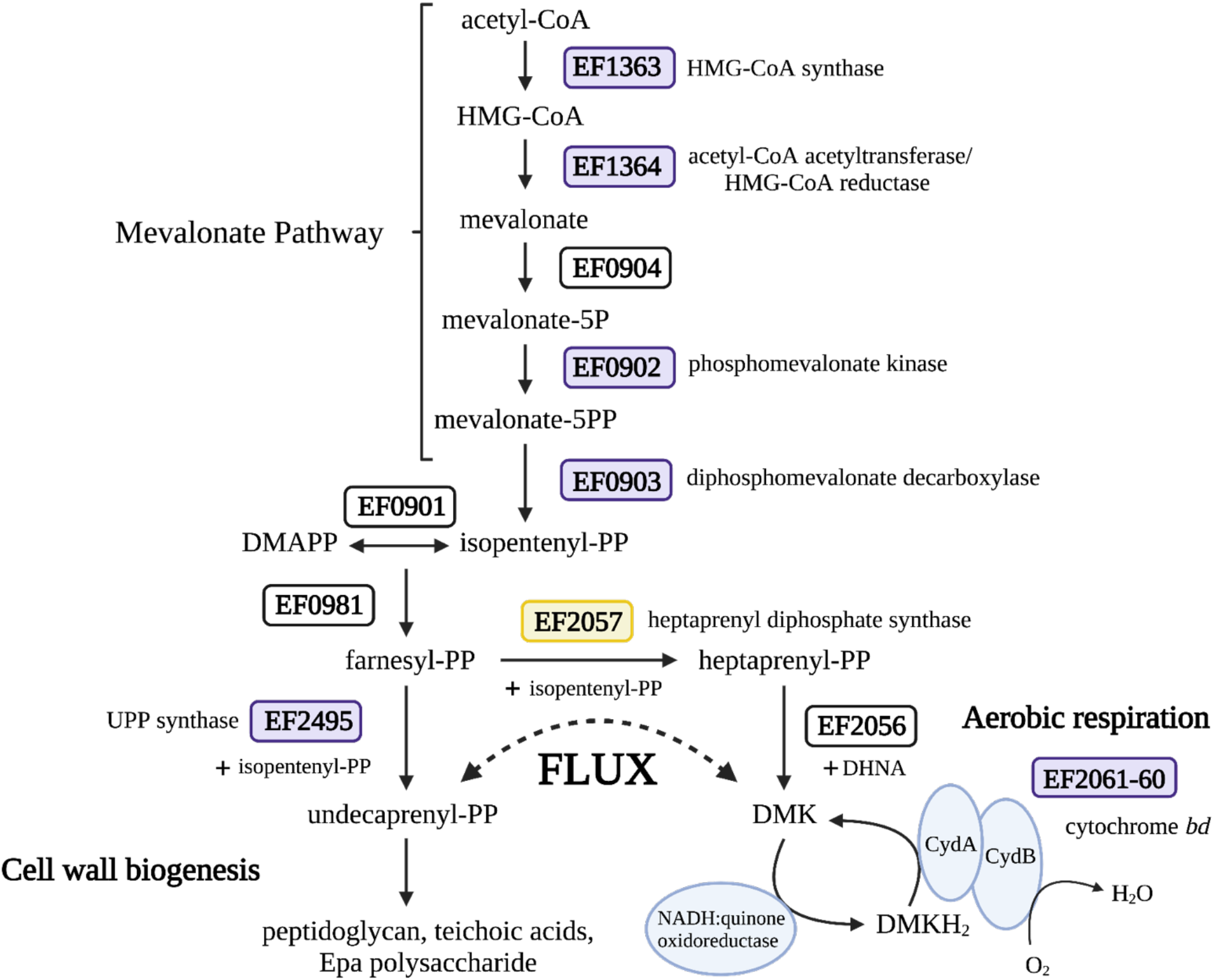
CroRS regulates isoprenoid flux between cell wall biogenesis and aerobic respiration to maintain cell wall homeostasis in response to antimicrobial stress. In wildtype *E. faecalis* CroRS confers tolerance to TXB-induced cell killing by regulating isoprenoid flux using a dual mechanism. In response to TXB challenge, we hypothesize CroRS simultaneously reduces the capacity for aerobic respiration, and thus the demand for isoprenoids/demethylmenaquinone (DMK), by decreasing the expression of cytochrome *bd* (purple box; CydA, CydB), while upregulating genes (purple boxes) involved in isoprenoid biosynthesis (i.e., mevalonate pathway and UPP synthase) and cell wall biogenesis (i.e., peptidoglycan, teichoic acids and Epa polysaccharide). In the absence of *croRS, E. faecalis* is susceptible to killing by TXB. However, proposed loss of function mutations in a heptaprenyl diphosphate synthase (yellow box; HppS) can partially rescue this phenotype. This supports our hypothesis that TXB tolerance can be conferred in *E. faecalis* through the control of isoprenoid flux from aerobic respiration to cell wall biogenesis.

Mutations in DNA-directed RNA polymerase subunits have previously been associated with antimicrobial susceptibility in enterococci (40, 41). Traditionally, mutations in *rpoB* have been associated with rifampicin resistance, however they, as well as mutations in *rpoC*, can also alter intrinsic cephalosporin and daptomycin resistance in *E. faecalis* (41–43). Uniquely, we identified a mutation in *rpoE* in the 3BS mutant, we hypothesize this may contribute to the recovery of tolerance to TXB and vancomycin in the Δ*croRS* background (**Table 2**).

EF2367 encodes an *N*-acetylmuramoyl-L-amidase, a putative autolysin. Autolysins play an essential role in normal cell wall turnover and their absence has been associated with increased penicillin tolerance in enterococci (44–46). In the RNA-Seq analyses above, we found that CroRS upregulated expression of teichoic acids, a known target of TXB and regulator of cell wall autolytic activity (47, 48). Therefore, it is tempting to speculate that in the absence of CroRS, teichoic acids are depleted and autolytic activity is dysregulated as a consequence. This may contribute to the increase in sensitivity to cell wall-acting antimicrobials in Δ*croRS*. In this scenario, loss of function mutations in putative autolysins, such as those observed in EF2367 in the 1BS and 5BS mutants, may reduce autolytic activity and subsequently confer the decrease in antimicrobial susceptibility observed in these mutants compared to the parental Δ*croRS* strain.

The phosphate ABC transporter, Pst, is a high-affinity phosphate transport system with a putative role in the uptake of phosphate under nutrient stress (49). In *E. faecalis* the *pst* operon consists of five *pst* genes (*pstS, C, A, BA* and *BB*) and *phoU*. In *E. faecium* a S199L amino acid substitution in PstBB confers protection against killing by vancomycin and chlorhexidine co-treatment (50), while in *S. pneumoniae* deletion of *pstB* resulted in a decrease in uptake of phosphate and reduced autolysis, likely due to an alteration in amidase activity (51). Perhaps then it is not a coincidence that we observed an amino acid substitution in in PstBB alongside a mutation in the putative autolysin, *N*-acetylmuramoyl-L-amidase, as both may be working together to rescue tolerance in the Δ*croRS* 5BS mutant by reducing autolytic activity (**Table 2**).

### Antimicrobial susceptibility and cell physiology are intricately linked through the cell envelope in *E. faecalis*

Alterations in cell physiology influence AMT in heterogenous sub-populations such as persisters (Fridman *et al.*, 2014; Brauner *et al.*, 2016; Levin-Reisman *et al.*, 2017). While mutations in *rpoE*, EF2367 and *pstBB* can be plausibly linked to the rescue of the Δ*croRS* strain, causation cannot yet be established as they do not occur on their own, unlike the mutations observed in *hppS*. Therefore, to determine the relationship between whole-population tolerance and cell physiology, the *E. faecalis* WT, Δ*croRS* and HppS-defective 4BS mutant strains were characterized in more detail. A TXB time-kill assay was carried out at 25 × MIC (50 *μ*g ml^-1^) over 24 h. After 24 h exposure, strong killing (6-log_10_ decrease) of the Δ*croRS* strain was observed, while the WT and 4BS mutant were reduced only 2 and 4-log_10_, respectively (**Figure 3A**). This showed that the 4BS mutant displayed an intermediate phenotype of the isogenic WT and Δ*croRS* parent strains. This intermediate phenotype was also observed in growth inhibition assays in the presence of various stressors, including perturbation of the cell envelope (glycine), osmotic stress (NaCl), and temperature stress (50 °C). In all conditions, sensitivity to cell stresses was drastically increased in the Δ*croRS* strain compared to the WT, while the 4BS mutant displayed partially restored growth (**Figure 3B – D**).

Given the impact of CroRS signalling on all layers of cell envelope biogenesis observed in the RNA-Seq analyses reported above, we hypothesized that the general sensitivity of the Δ*croRS* strain to conditions that are likely to affect cell envelope integrity may be due to physical changes in this cellular structure. We therefore assessed the cell size and cell envelope thickness using transmission electron microscopy. The first observation was that the Δ*croRS* strain was significantly larger in cell size and had a thinner cell envelope than the WT (**Figure 4A, B and D**). This was consistent with our hypothesis that loss of CroRS impacts cell envelope integrity. Interestingly, the 4BS isolate showed a significant reduction in cell size compared to the Δ*croRS* strain, and was comparable to the WT (**Figure 4A**), and it showed a significant increase in cell envelope thickness compared to Δ*croRS* (**Figure 4B**). However, the latter was not fully restored to WT, perhaps explaining the intermediate physiology observed in the stress analyses (**Figure 3 and Figure 4**).

The transcriptomic and phenotypic evidence presented here suggests that the Δ*croRS* strain is in a cell envelope-stressed state even in the absence of antimicrobials (**Figure 3** and **Figure 4**). If our hypothesis is correct that the deletion of *croRS* compromises cell envelope integrity, which can be rescued by a truncation of HppS, we should be able to detect this using a reporter assay for cell envelope stress. LiaFSR is another two-component system in *E. faecalis* which is well-known to respond to the perturbation of cell envelope integrity (10). To test for cell envelope stress, a transcriptional reporter of its target promoter, P_*liaX*_, and the *lacZ* gene was developed and used to transform the Δ*croRS* strain alongside the WT and 4BS suppressor mutant (54). In the absence of antimicrobial stress, the WT strain displayed very low P*_liaX_-lacZ* activity as expected, while the Δ*croRS* strain showed very high constitutive activity (>150-fold increase) (**Figure 4C**), a clear indicator that this strain is indeed in a state of constant cell envelope stress due to the loss of CroRS. Interestingly, the single stop codon mutation in *hppS* carried by 4BS was sufficient to reduce this stress to near WT levels (**Figure 4C**). This lends weight to our hypothesis that the main phenotype(s) of the Δ*croRS* mutant are due to changes in the mevalonate pathway and aberrant flow of isoprenoids. As UPP is the shuttle molecule on which peptidoglycan, teichoic acids and cell wall polysaccharides are moved to the extracellular face of the cytoplasmic membrane, it is plausible that shortages in UPP provision would cause defects in cell wall and wider cell envelope biogenesis.

## Conclusion

### Isoprenoid flux plays a key role in CroRS-mediated antimicrobial tolerance in *E. faecalis*

HppS is a component of isoprenoid metabolism, which starts with the MVA pathway and leads to cell wall synthesis (UPP) and aerobic respiration (DMK) (**Figure 5**), with HppS the first step of the latter branch. Suppressor mutations of a putative HppS were here shown to rescue AMT in the Δ*croRS* background. We propose loss of function mutations in HppS allow rerouting of FPP and IPP to UPP and consequently cell wall biosynthesis, to rescue AMT in the Δ*croRS* background. Interestingly, mutations in the MVA pathway in *E. faecalis, S. aureus*, and *S. pneumoniae* have been shown to play a critical role in peptidoglycan biosynthesis, with suppressor mutations in *S. pneumoniae* decreasing peptidoglycan precursors and resensitizing the bacterium to amoxicillin (55, 56). In addition, shortages in peptidoglycan precursors were shown to cause elongation of cells, reminiscent of the enlargened cells we observe in Δ*croRS* (55) (**Figure 4D**).

Based on the data presented in this study, we propose a model where CroRS acts as a gate-keeper between the two branches of isoprenoid biosynthesis, controlling the flux of isoprenoids needed for cell wall synthesis and respiration, to maintain cell wall homeostasis (**Figure 5**). When attacked by antimicrobials in WT *E. faecalis*, CroRS can increase MVA pathway flux to ensure sufficient UPP production so cell wall synthesis is maintained. But as a consequence, this also increases production of respiratory players, as the flow of isoprenoids would follow enzyme kinetics and not active decision making, and will likely lead to increased production of DMK and derived compounds. This could pose a risk of increasing ROS and ROS-derived cell damage. However, CroRS counteracts this by actively downregulating cytochrome *bd* (**Figure 5**). The resulting reduction in respiratory activity may also reduce the demand for isoprenoids for the DMK branch of isoprenoid metabolism, further increasing precursor availability for UPP synthesis. Conversely, in the Δ*croRS* strain, where this control of isoprenoid flux is missing, cell envelope integrity and regulatory control of cytochrome *bd* expression are lost. This could explain the loss of tolerance in the Δ*croRS* strain, with susceptibility to both modes of antimicrobial-induced killing, i.e., blocking of cell wall synthesis and production of ROS, increased. This model also explains the suppressive effect of loss of function mutations in *hppS*: blocking the DMK branch of isoprenoid metabolism would increase the availability of isoprenoids for UPP synthesis to aid in cell wall biosynthesis, while potentially also reducing respiratory activity and ROS production (**Figure 5**). Interestingly, the *hppS* mutations corrected the growth defect and survival (i.e., tolerance) against TXB and vancomycin of the Δ*croRS* mutant, but could not counteract the inhibitory effects of the antibiotics on cell proliferation (i.e. resistance). This clearly shows that the mechanims of growth inhibition and killing by these antibiotics, and by extension the mechanisms of resistance and tolerance, are functionally separate.

Our study has revealed key insights into the regulatory role CroRS plays in controlling the cellular response against antimicrobial-induced killing and explains its strong impact on AMT in *E. faecalis*. Future work will endevour to further deconstruct the relationship between tolerance and resistance and to identify the molecular mechanisms deployed by *E. faecalis* against inhibition of growth proliferation.

## Methods

### Bacterial growth/growth curves

All *E. faecalis* strains were routinely grown in BHI broth and agar overnight at 37°C with no aeration unless otherwise stated. All *Escherichia coli* strains were routinely grown in LB broth and agar at 37°C (200 r.p.m), unless otherwise stated. Cultures for RNA sequencing and optimization were grown as previously described (12). Growth was determined by optical density at 600 nm (OD_600_). TXB stocks were made with dimethyl sulfoxide (DMSO) and stored at - 20°C. All bacterial strains used in this study are listed in **Table S6.**

### Antimicrobial susceptibility assays

Minimum inhibitory (MIC) and bactericidal (MBC) concentrations were determined as previously described (12). Time-dependent kill assays were carried out to determine cell death kinetics over time as previously described (12).

### RNA extraction and preparation of RNA samples for sequence analysis

*E. faecalis* JH2-2 and *DcroRS* were grown in biological triplicate to mid-exponential phase (OD_600_ of 0.5) at 37°C, 130 r.p.m. Cultures were subsequently split to produce two sets of biological triplicates. One set was challenged with 0.5 *μ*g ml^-1^ (0.25 × MIC) of TXB, while the other set were untreated controls. Cultures were grown for a further 1 h and harvested for RNA extraction. Total RNA was isolated using TRIzol-chloroform extraction as previously described (12). RNA samples were purified using a Zymo RNA clean and concentrate kit as per the manufacturer’s instructures. RNA concentration and integrity (RIN >8) was determined by bioanalyzer.

### RNA sequencing and gene expression analysis

#### (i) cDNA library preparation and sequencing of the E. faecalis JH2-2 and ΔcroRS transcriptomes

RNA libraries were prepped using the Zymo-Seq RiboFree Total RNA-Seq Library Kit. Sequencing was complete using an Illumina MiSeq (v3) system generating 150 bp paired-end reads.

#### (ii) Analysis of RNA sequencing data

Adapter sequences were removed from raw fastq files using bbduk. Reads were aligned to the *E. faecalis* JH2-2 genome (GenBank accession numbers NZ_KI1518257.1 and NZ_KI1518256.1) using Bowtie2. Statistical and principle-component analyses were performed using the Bioconductor DESeq2 package. Parameters considered during analysis were the fold change (≥ 1.0-fold log_2_), the mean number of reads (>50), and the adjusted *P* value (P_*adj*_ <0.1). Genes were also annotated with the *E. faecalis* V583 reference (GenBank accession number NC_004668.1) gene homolog and ontology using KEGGRest. Gene annotations and ontology assignments were complemented with NCBI BLAST and literature searches when necessary.

#### (iii) Hypergeometric testing to determine gene ontology enrichment

Following assignment of differentially expressed genes to KEGG pathways, a hypergeometric test was performed by comparing the number of genes within the regulon to the total number of genes within the respective *E. faecalis* V583 KEGG pathway. Pathways with a *P* value <0.05 were deemed significantly enriched.

### Serial passaging of ΔcroRS for wild-type growth

To isolate supressor mutants with improved fitness compareed to the *croRS* deletion strain, we utilised a serial passaging method. Five independent overnight cultures were inoculated 1:1000 into 10 ml BHI medium and incubated at 37°C with no aeration. Cultures were passaged every 48 h under the same conditions. Cells were serially diluted and plated every three passages onto BHI and bile-esculin medium to ensure no contamination had occured. The experiment was concluded upon visible confirmation of improved fitness, assessed as increased turbidity of overnight cultures, which appeared within 10 – 14 days.

### Isolation of vancomycin-tolerant ΔcroRS mutants

*E. faecalis* Δ*croRS* were grown to an OD_600_ of 0.5 and diluted 1/100 (0.005) and 1/1000 (0.0005) and exposed to vancomycin at 50 × MIC for 24 h. 1ml of cells were pelleted (12,000 × *g* for 1 min) and washed twice in equivolume 1 × PBS. Cells were resuspended in 100 μl 1 × PBS and plated on BHI agar containing no antimicrobial for incubation overnight at 37 °C. Plates were observed for colonies the following day and screened for recovery of vancomycin tolerance using standard MIC and MBC assays.

### General stress testing

#### Temperature stress

In biological triplicate, overnight cultures were normalised to an OD_600_ of 0.5 and inoculated 1/10 into fresh BHI broth to a final OD_600_ 0.05. Cells were grown at 37°C and 50°C overnight with no aeration. Following incubation, cell growth was measured by optical density (600 nm) and calculated as a percentage comparison of before and after temperature challenge.

#### Osmotic (NaCl) stress

In biological triplicate, overnight cultures were pelleted by centrifugation (10,000 × *g* for 5 min) and washed in 1 × phosphate saline buffer. Cells were resuspended in BHI + NaCl (6.5% or 13%) to an OD_600_ of 0.05 and grown overnight at 37°C with no aeration. Following incubation, cell growth was measured by optical density (600 nm) and calculated as a percentage comparison of before and after osmotic challenge.

#### Cell wall (glycine) stress

In biological triplicate, overnight cultures were diluted 1/200 in BHI + glycine at concentrations of 0, 3, 5, 6, 7, 8, and 9% (w/v). Cultures were grown overnight at 37°C with no aeration. Cell growth was measured by optical density (600 nm) and calulated as a percentage comparison to the untreated control (0% glycine).

### Transmission electron microscopy of E. faecalis WT, ΔcroRS and 4BS strains

*E. faecalis* strains were grown to mid-exponential phase (OD_600_ of 0.5, 1 × 10^8^ CFU ml^-1^) in fresh BHI broth and harvested by centrifugation (3000 r.p.m, 4°C, 6 min). Cell pellets were resuspended in a primary fixative solution (5% glutaraldehyde in 0.1 M cacodylate buffer) for 30 min at room temperature. Cells were subsequently washed 3 × in 0.1 M cacodylate buffer and incubated with a secondary fixative solution (1% osmium and 1% potassium ferricyanide in 0.1M cacodylate buffer) for 30 min. Following application of the secondary fixative, cells were washed 3 × in 0.1 M cacodylate buffer and stored in the refrigerator for further processing. Fixed cells were washed with ddH_2_O and dehydrated with ethanol at progressively increasing concentrations: 30% ethanol for 5 min, 50% ethanol for 5 min, 70% ethanol for 5 min, and 95% ethanol for 5 min, followed by three treatments with 100% ethanol for 10, 15, and 20 min, respectively. The dehydration process was completed by 2 × 15 min treatments with propylene oxide. Finally, cells were treated for 2 h with a 1:1 mixture of resin and propylene oxide, then left in resin overnight. Following this incubation, cells were embedded in fresh EmBed 812 resin and polymerized at 60 °C for 48 h. Ultrathin sectioning (85 nm) was carried out using a diamond knife on a Leica EM UC7 ultramicrotome and collected on formvar-coated copper grids. Sections were stained with uranyl acetate and lead citrate and investigated using a Philips CM100 BioTWIN transmission electron microscope (Philips/FEI Corporation, Eindhoven, Holland) fitted with a MegaView lll digital camera (Soft Imaging System GmbH, Münster, Germany).

### Construction of the liaRS promoter-lacZ plasmid

The *liaX* transcriptional promoter fusion to *lacZ* in *E. faecalis* was constructed in the vector pTCVlac (57). The *liaX* promoter fragment was amplified with the primers 5’-AATTT<u>GAATTC</u>GGATGATCGTACTAATG-3’ and 5’-AATTT<u>GGATCC</u>CTTTCATGGATATTGC-3’. The fragment was then inserted into pTCVlac via the EcoRI and BamHI restriction sites. The resulting plasmid PliaX_pTCVlac was transformed into electrocompetent *E. faecalis* WT, Δ*croRS* and 4BS strains by electroporation (**Table S6**).

### β-galactosidase assays

To quantitatively assess induction of the PliaX-*lacZ* reporter construct in *E. faecalis*, cells were grown to mid-exponential phase (OD_600_ 0.4 – 0.5) in BHI medium. Cells were harvested via centrifugation and stored at −20°C. *β*-galactosidase activity was assayed in permeabilised cells and expressed in Miller units (MU). For this, cells were resuspended and normalised to an OD_600_ of 0.5 in Z-buffer. From this cells were diluted 1/5 and 1/2.5 in Z-buffer and lysed by vortexing with 0.1% SDS (w/v) and chloroform. Cells were left to rest at room temperature for 5 – 10 min. Reactions were initiated by the addition of ONPG (*o*-nitrophenyl-*β*-D-galactopyranoside; 4 mg ml^-1^ in Z buffer) and monitored for yellow colouration. Reactions were monitored for colour change for a maximum of 20 min. Upon colour change, reactions were stopped by adding 1M sodium carbonate and reaction time noted. Absorbance at 420 nm (A420) was the read and MU calculated using the following equation.

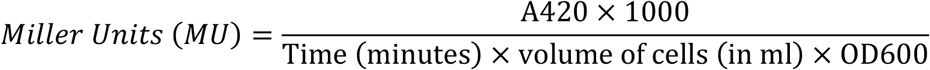

## Acknowledgements

All authors acknowledge funding support from the Health Research Council (HRC) (NZ), the Maurice Wilkins Centre for Molecular Biodiscovery (NZ), and the University of Otago Research Grant (NZ). F. O. Todd Rose was supported by a University of Otago PhD Scholarship (NZ), and S. Morris was supported by a GW4 BioMed Medical Research Council (MRC) Doctoral Training Partnership Scholarship (UK). The authors would like to thank Dallas Hughes from Novobiotic for the kind gift of teixobactin. The authors would also like to thank Fatima Esperanca Jorge and Richard Easingwood at the Otago Micro and Nano Imaging Unit, University of Otago for their assisatnce in the preparation and collection of the transmission electron microscopy images.

## Supplementary Figures

**Figure S1.**
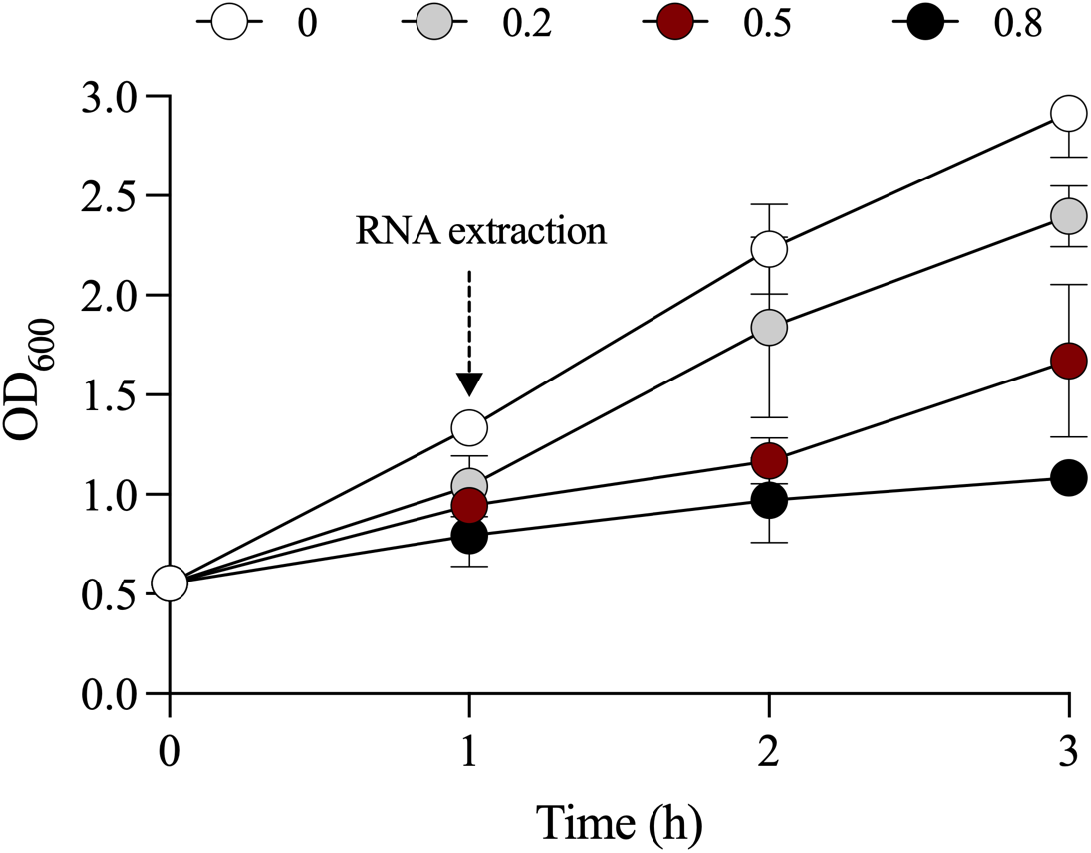
Optimisation of teixobactin concentration for RNA sequencing. *E. faecalis* Δ*croRS* was grown to mid-exponential phase in BHI broth and challenged with TXB at concentrations of 0, 0.2, 0.5, and 0.8 *μ*g ml^-1^ for 3 h. Growth was measured by OD_600_ at 1, 2, and 3 h post-challenge. Following optimisation, RNA was subsequently extracted from cultures challenged with and without 0.5 *μ*g ml^-1^ of TXB at 1 h post-challenge. Data is representative of the mean of biological triplicate ± SD.

**Figure S2.**
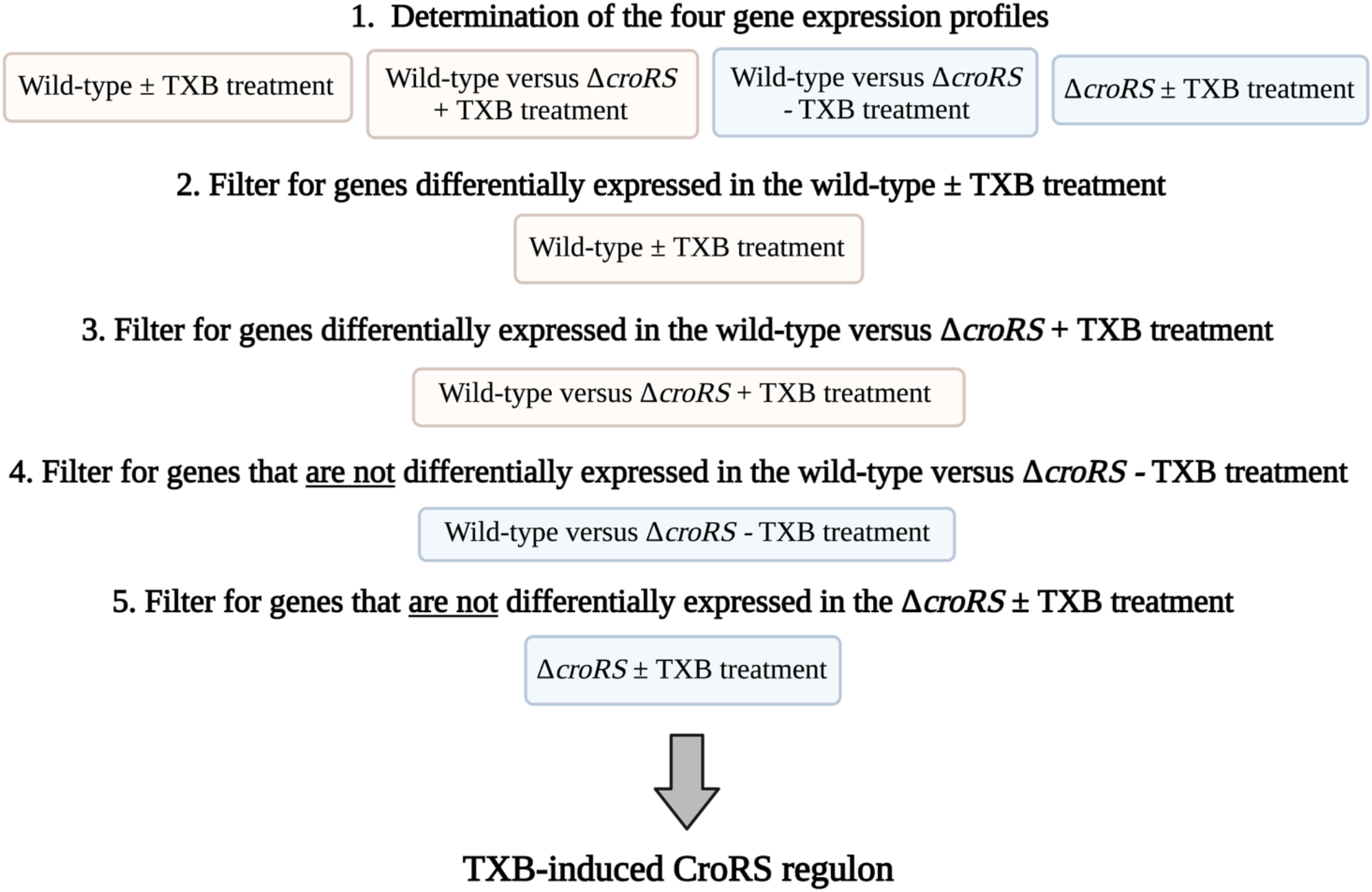
Determing the TXB-induced CroRS regulon. To determine the teixobactin (TXB)-induced CroRS regulon we first determined the four gene expression profiles. Each profile was subsequently filtered using the criteria listed above (Steps 2 – 5) to reach a final set of genes which represented the TXB-induced CroRS regulon.

**Figure S3.**
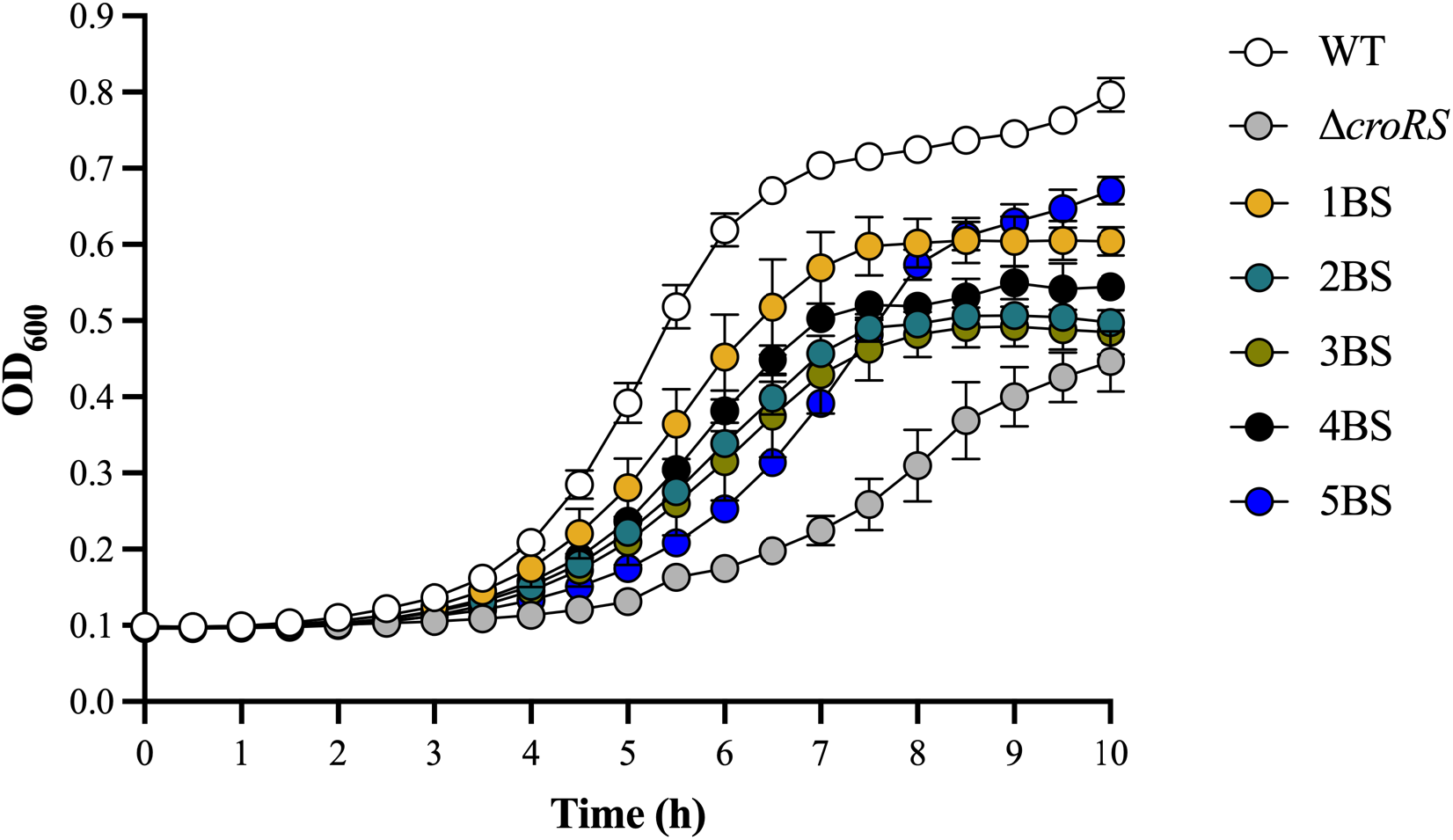
Growth curve of the evolved Δ*croRS* mutants 1BS – 5BS alongside the isogenic Δ*croRS* mutant and WT (JH2-2). Overnight cultures of *E. faecalis* JH2-2, Δ*croRS*, and Δ*croRS*-1BS to 5BS were diluted in fresh BHI and inoculated to an OD_600_ of 0.01 in 96-well plates. Cells were incubated at 37°C with no aeration and growth was monitored as optical density (OD_600_) with readings taken every 30 min for 10 h. Results are presented in biological triplicate ± SD.

**Figure S4.**
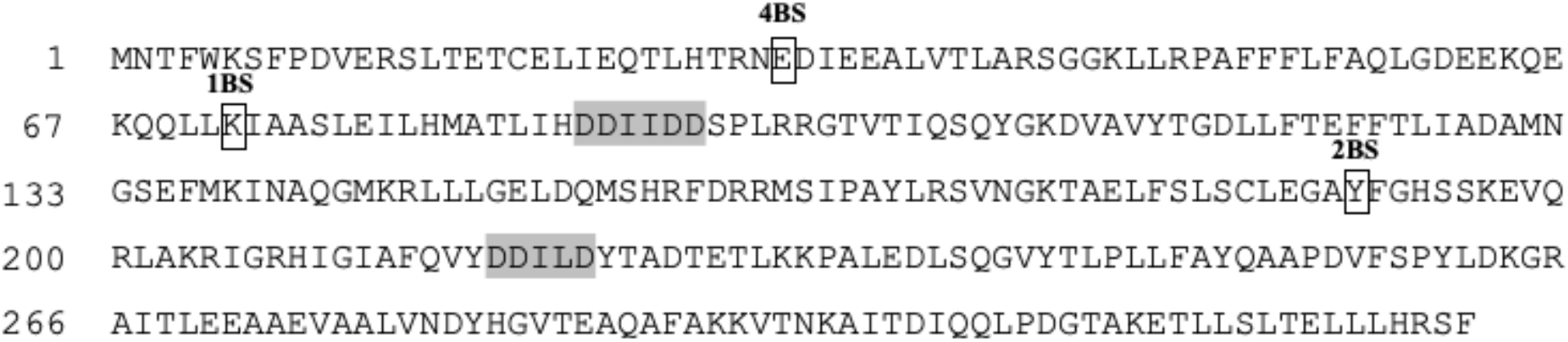
The amino acid sequence of the putative heptaprenyl diphosphate synthase identifying mutations in the gain of tolerance Δ*croRS* mutants. Two aspartate-rich motifs (DDXXD) are essential and highly conserved among (E)-prenyl diphosphate synthases (grey). The relative positions of the mutations found within the strains van1, 1BS, 2BS and 4BS are indicated and the amino acid containing the mutation is box. Mutants 2BS and 4BS result in a truncation of the protein at positions Y189 and E31 respectively, due to the introduction of a STOP codon, while in the 1BS mutant, a deletion of 9 bp (216_225) results in a frameshift at K72.

